# Epicardial reservoir-enabled multidose delivery of exogenous FSTL1 leads to improved cardiac function, healing, and angiogenesis

**DOI:** 10.1101/2022.11.02.513725

**Authors:** Claudia E. Varela, David S. Monahan, Shahrin Islam, William Whyte, Jean Bonnemain, Souen Ngoy, Sudeshna Fisch, Garry P. Duffy, Ellen T. Roche

## Abstract

Epicardial delivery of human follistatin-like 1 protein (FSTL1) induces significant cardiac benefit following a myocardial infarction (MI). However, the optimal dosing regimen for maximal therapeutic benefit has not yet been elucidated. To investigate the impact of multiple FSTL1 doses, without the confounding effects of multiple surgical procedures for multidose delivery, alternative delivery strategies are needed. Here, we use an epicardial reservoir that allows non-invasive delivery of additional doses after implantation to investigate the impact of single, double, and triple FSTL1 dose regimens in a rat model of MI. Multidose delivery of FSTL1 improves ejection fraction (3 doses), fractional shortening (1, 2 and 3 doses), and chamber stiffness (2 doses) 28 days after MI. Histologically, multiple FSTL1 doses increase ventricular wall thickness (2 and 3 doses) and reduce infarct size (1, 2, and 3 doses). We also demonstrate a dose-dependent increase in blood vessel number and density in the infarct zone, with three FSTL1 doses leading to the highest improvements. This study shows that multidose delivery of FSTL1 improves cardiac function, healing, and angiogenesis following MI. The epicardial delivery platform used here may be essential in optimizing dosing regimens of various bioagent combinations for a range of clinical indications.

## 1. Introduction

Over 1 million people suffer from a myocardial infarction (MI) in the United States every year.^[1]^ A myocardial infarction (MI), or heart attack, occurs when blood flow to a region of myocardium is interrupted, depriving cardiomyocytes of oxygen, leading to a loss of contraction, necrosis of the infarcted region, and subsequent scar formation. Often, if the infarct is large, the ventricle undergoes adverse tissue remodeling, and can ultimately lead to heart failure (HF)—a complex clinical syndrome where the heart cannot pump enough blood to meet the body’s metabolic demands. Current clinical management of MI focuses on limiting the amount of muscle damage by reopening blocked arteries early after MI (e.g., thrombolytics, angioplasty, coronary stenting, coronary artery bypass) or on managing HF via medication, lifestyle changes, mechanical circulatory support, and organ transplantation. Despite these treatments, HF remains a prominent cause of morbidity and mortality in MI survivors.^[1]^ To date there is an unmet clinical need for a therapeutic strategy that promotes early recovery of cardiac muscle/function after MI. Pre-clinical evidence indicates that delivering bioagents, such as stem cells, growth factors, proteins, and drugs, directly to the infarcted heart can lead to an early improvement in cardiac recovery.^[2–16]^ However, studies that optimize bioagent timing and dosing are required to maximize their therapeutic effect and promote clinical translation.

Most rodent studies have delivered a limited amount of bioagent (i.e., a single dose) to the heart during the MI creation surgery using direct myocardial injections or coupling a biomaterial carrier to the heart surface.^[2–16]^ Using the latter delivery method, human follistatin-like 1 protein (FSTL1) was shown to promote cardiac regeneration and repair following MI.^[17]^ Using mouse and swine models of MI, Wei & Serpooshan et al.^[17]^ demonstrated that a single dose of exogenous FSTL1 delivered through a collagen patch sutured onto the epicardial surface leads to improvements in animal survival, cardiac function, healing, and cardiomyocyte regeneration. Although interventions with FSTL1 are very amenable to clinical translation, the FSTL1 dosing regimen that yields maximal therapeutic benefits remains unknown. Interestingly, Magadum et al. reported no cardiac benefit from repeated FSTL1 induction via injection of an mRNA construct.^[18]^ However, they hypothesized that factors associated with the secondary surgical procedure, such as the inflammatory environment and physiological injury to the heart, may have confounded their results. Other studies have found that multiple doses of surgically delivered bioagents results in cumulative therapeutic benefits.^[19,20]^

Enabling non-invasive delivery of additional bioagent doses is required to dissociate their therapeutic benefit from the potentially detrimental effects of additional invasive interventions. To this end, we used an epicardial delivery platform^[21]^ that enables localized, non-invasive replenishment of bioagents to deliver additional FSTL1 doses. The delivery platform consists of an epicardial reservoir implanted at the time of surgical MI creation and is connected to a subcutaneous injection port via a catheter line which enables non-invasive delivery of additional bioagent doses. With this platform, we achieved the goal of this study—characterizing the impact of a single, double, and triple dose regimen of FSTL1 on cardiac function, scarring and angiogenesis in a rat model of MI.

## 2. Materials and Methods

### Animal study

Animal procedures were reviewed and approved according to ethical regulations by the Institutional Animal Care and Use Committees at Brigham and Women’s Hospital and the Massachusetts Institute of Technology. For this 28-day study **(Figure 1a)**, female Sprague Dawley rats (225-250 g) were assigned to the following groups: 1) sham device (epicardial device with PBS-loaded gelatin sponge), 2) single FSTL1 dose (via FSTL1-loaded gelatin sponge), 3) 2 doses of FSTL1 (FSTL1-loaded gelatin sponge and 1 refill via epicardial device), and 4) 3 doses of FSTL1 (FSTL1-loaded gelatin sponge and 2 refills via epicardial device). Functional and histological findings in rats that received no treatment (MI only) were measured using the same methods and personnel and reported previously.^[21]^

**Figure 1.**
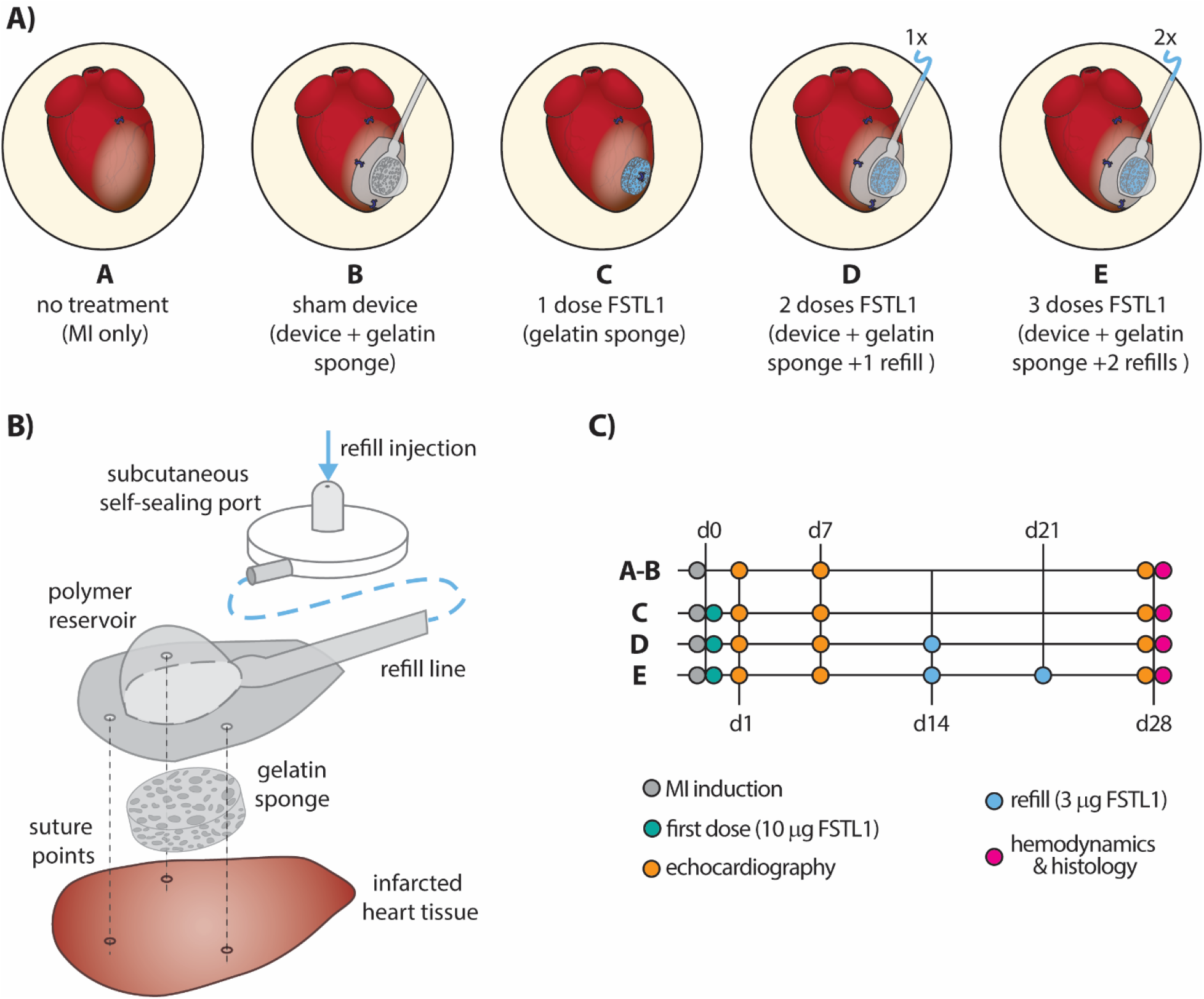
Animal study overview and epicardial reservoir design. **A)** Description of animal study groups. **B)** Schematic of sutured epicardial reservoir (exploded view) and trajectory of dose refill after subcutaneous port injection. **C)** Timeline for animal study, including corresponding assessments and FSTL1 dosing regimens.

### Epicardial device fabrication

All epicardial devices were manufactured as previously described. ^[21]^ A thermoplastic polyurethane (TPU) sheet (HTM 8001-M polyether film, 0.003” thick, American Polyfilm, Inc) was formed into a hemispherical reservoir using a vacuum thermal former (Yescom Dental Vacuum Former, Generic). The neck of the reservoir was heat bonded to a 10 cm, 3 Fr thermoplastic polyurethane catheter (Micro-Renathane 0.037” × 0.023”, Braintree scientific) using a heat transfer machine (Heat Transfer Machine QXAi, Powerpress). Three holes were created around the perimeter of the reservoir to guide suture placement during implantation. The assembled device was sterilized using a low temperature ethylene oxide cycle.

### Surgical procedure

MI creation, implantation of the gelatin sponge and/or epicardial device were performed as previously described. ^[21]^ Briefly, rats that received a device implantation were anaesthetized (1–3% isoflurane in O_2_), shaved at the dorsal (between the shoulder blades) and ventral (left side of the chest) surgical sites, and injected with a regional nerve blocker (lidocaine/bupivacaine) and a pre-operative analgesic (buprenorphine, 0.05 mg/kg) subcutaneously. Endotracheal intubation was performed, and subcutaneous incisions were made at the dorsal and ventral surgical sites. No dorsal incision was performed in groups where no epicardial device was implanted. The device refill line was tunneled subcutaneously from the dorsal to the ventral incision before MI creation. A thoracotomy was performed between the fourth and fifth intercostal space and the pericardium was removed using forceps. A MI was created by permanently ligating the left anterior descending (LAD) coronary artery with a suture (6-0 or 7-0 prolene) approximately one third of the way from the base to the apex of the heart. After MI induction, a gelatin sponge was either directly sutured onto the heart (single suture, 7-0 prolene) or placed inside the device reservoir for subsequent attachment. The reservoir was sutured (7-0 prolene) to the surface of the heart at 3 previously marked points **(Figure 1b).** After gelatin sponge or reservoir attachment, the thoracotomy and ventral incision were closed (layer-by-layer, 4-0 vicryl). In rats with an epicardial device, a vascular access button, or self-sealing port (VAB62BS/22, Instech Laboratories), was then connected to the end of the device refill line and placed subcutaneously between the shoulder blades of the rat. The port was secured to the underlying fascia using at least two interrupted sutures (4-0 vicryl) prior to skin closure. Animals were ventilated with 100% oxygen on a heated pad until autonomous breathing was regained. 3 ml of warm saline was administered subcutaneously, and buprenorphine (0.05 mg/kg in 50 *μ*l IP) was given every 12 h for three days post-operatively.

### FSTL1 delivery

The initial dose of FSTL1 was delivered to the treatment groups through a FSTL1-loaded gelatin sponge. A sterile 4-mm biopsy punch was used to cut a core of commercially available sterile gelatin sponge (Gelfoam®, or SurgiFoam®) inside a biological hood. The core was soaked for at least 30 min prior to implantation in a solution containing 10 *μ*g of FSTL1 (Aviscera Bioscience, suspended in 30 or 100 *μ*l of sterile PBS). Forceps were used to gently tap the gelatin sponge core to promote absorption of the FSTL1 solution. FSTL1-loaded gelatin sponges were secured directly onto the infarct region of rats in the single dose group (without the epicardial device) or placed inside the device reservoir prior to suturing the device to the heart in the double and triple dose groups. A PBS solution was used to soak the gelatin sponge cores placed inside the devices implanted into the sham device group.

The double and triple dose groups received either 1 or 2 FSTL1 refills, respectively. The first FSTL1 refill (second dose) occurred at day 14 post-MI while the second refill (third dose) occurred at day 21 post-MI **(Figure 1c).** For each refill, the rat was anesthetized (1–3% isoflurane in O_2_) and negative pressure was applied to clear the refill line of the device using an empty syringe connected to the self-sealing port. Then, 3*μ*g of FSTL1 (Aviscera Biosciences, suspended in 30 *μ*l of sterile PBS) were injected through the port into the refill line followed by 20 *μ*l of PBS to clear the refill line from any FSTL1 and ensure the FSTL1 reached the epicardial reservoir.

### Echocardiography

Echocardiography was conducted at days 1, 7 and, 28 for all groups **(Figure 1c)**. Data was acquired using either a Vevo 2100 or 3100 Ultrasound machine and a MS200 transducer probe (9-18Mhz). End systolic volumes (ESV), end diastolic volumes (EDV) and ejection fraction (EF) measurements were obtained from B-mode, parasternal long axis images. Fractional shortening (FS) measurements were obtained using an M-Mode parasternal short axis view of the heart, at mid ventricular level, with papillary muscles evident. Data was analyzed using the Visualsonics software (VevoLAB). The imaging datasets were analyzed and reviewed by 2 experienced ultrasound readers blinded to the study groups.

### Hemodynamic evaluation

At day 28, invasive hemodynamic recordings were performed as previously described.^[21,22]^ Briefly, a sternotomy was performed, a 5-0 silk suture snare was placed around the inferior vena cava (IVC), and a pressure-volume (PV) catheter (Millar SPR-838) was inserted into the left ventricle via an apical stick. After baseline recordings, PV relationships over a range of preloads were recorded as the IVC was temporarily occluded using the placed snare. LabChart Pro 8 (AD Instruments) was used for all PV data analysis. The volume signal was calibrated by inputting ESV and EDV obtained from echocardiography measurements. End diastolic PV relationships (EDPVRs) were obtained by performing an exponential curve fit of ~10 PV loops recorded during an IVC occlusion with LabChart’s PV loop module (*Ped*=*α*\**exp (β*\**Ved)*, where *Ped* and *Ved* respectively correspond to end diastolic pressure (mmHg) and volume (*μ*l), and *α* and *β* are fit parameters). The slope (*β*) of 1-4 EDPVRs was averaged to derive each animal’s chamber stiffness value.

### Histology

At the terminal point of the study, rats were anesthetized, and their hearts were arrested in diastole by direct injection of 2-3 ml of 1% *KCl* solution and fixed in a 10% neutral buffered formalin solution for 24 h. Explanted hearts were sliced into three transverse sections (apex, mid-ventricle, and base). Heart slices were placed into cassettes and processed using a tissue processor through increasing grades of ethanol to paraffin wax. Samples were then wax embedded and sectioned into 5-*μ*m slices using a microtome and mounted onto glass slides for histochemistry and immunohistochemistry. Hematoxylin and eosin (H&E) and Masson’s Trichrome staining was performed using previously established protocols.^[23]^ CD31 immunohistochemistry was performed to stain for blood vessels. Briefly, samples were dewaxed and rehydrated through decreasing grades of ethanol, and heat mediated antigen retrieval was performed using sodium citrate with 0.01% tween. Next samples were blocked using peroxidase, incubated with primary CD-31 antibody for one hour at room temperature (ab182981), and stained with 3,3 diaminobenzidine (DAB) using the Abcam staining kit (ab236469).

### Stereology

All quantitative analysis was performed using ImageJ from a blinded counter. For all measurements, test fields were gathered and a point grid with an adjusted number of test points was projected onto each test field. H&E and Masson’s trichrome stained samples were used to measure left ventricular thickness. A perpendicular line was drawn from each point where the grid intersected the inner aspect of the left ventricular wall to the outer aspect and the length was measured using the measure function with a defined scale bar. Masson’s trichrome stained hearts were used to determine the extent of myocardial scarring. The area fraction of scar tissue was estimated by applying a grid to the tissue and counting the number points overlaying scar tissue on the left ventricle which was divided by the number of points overlaying the left ventricle and expressed as a percentage. It was important that less dense fibrous capsule scar was removed during this measurement. Stereological measurements of blood vessels were performed as previously shown.^[23,24]^ Briefly, ten non-overlapping images were taken from the infarct zone, border zone, and adjacent myocardium. The number of blood vessels was quantified using a systematic randomized sampling approach with an unbiased counting frame. The number of blood vessels per area was determined and the length density (2* the number per area) and radial diffusion distance were determined (1/√π * Length density) for each heart. For each type of analysis, 2-4 sections were analyzed from different regions of the heart.

### Statistics

Statistical analysis was performed using GraphPad Prism. All graphs are expressed as a mean (± standard deviation) with means for individual animal shown as points. Normality tests were performed using a Shapiro-Wilks test and either a one-way ANOVA with post hoc Tukey’s test, or a 2-way ANOVA (Mixed model) with post Dunnet’s test were performed to test for significance. Differences between groups were considered statistically significant to each other if p<0.05.

## 3. Results and Discussion

### 3.1. Multidose FSTL1 regimes lead to improved cardiac function

We compared how left ventricular ejection fraction (EF) **(Figure 2a)** and fractional shortening (FS) **(Figure 2b)**, both echocardiography-derived cardiac function metrics, were impacted by the single, double, and triple FSTL1 dosing regimens over the course of the 28-day study. By day 7 after MI, FSTL1-treated animals had significantly better cardiac function than the MI only group after only one dose of FSTL1 had been delivered. Specifically, the single and double dose groups had higher EF **(Figure 2a)** and the double and triple dose groups had higher FS **(Figure 2b)** when compared to the MI only group. By day 28 after MI, only animals treated with 3 doses of FSTL1 had improved EF **(Figure 2c)** when compared to untreated controls. Although the FSTL1-treated and sham device groups had better FS than MI only animals at day 28, 3 doses of FSTL1 seem to yield the most FS improvement **(Figure 2b,d).** To our knowledge, this is the first study to demonstrate that FSTL1 delivery leads to improved cardiac function in a rat model of MI using a gelatin sponge biomaterial carrier.

**Figure 2.**
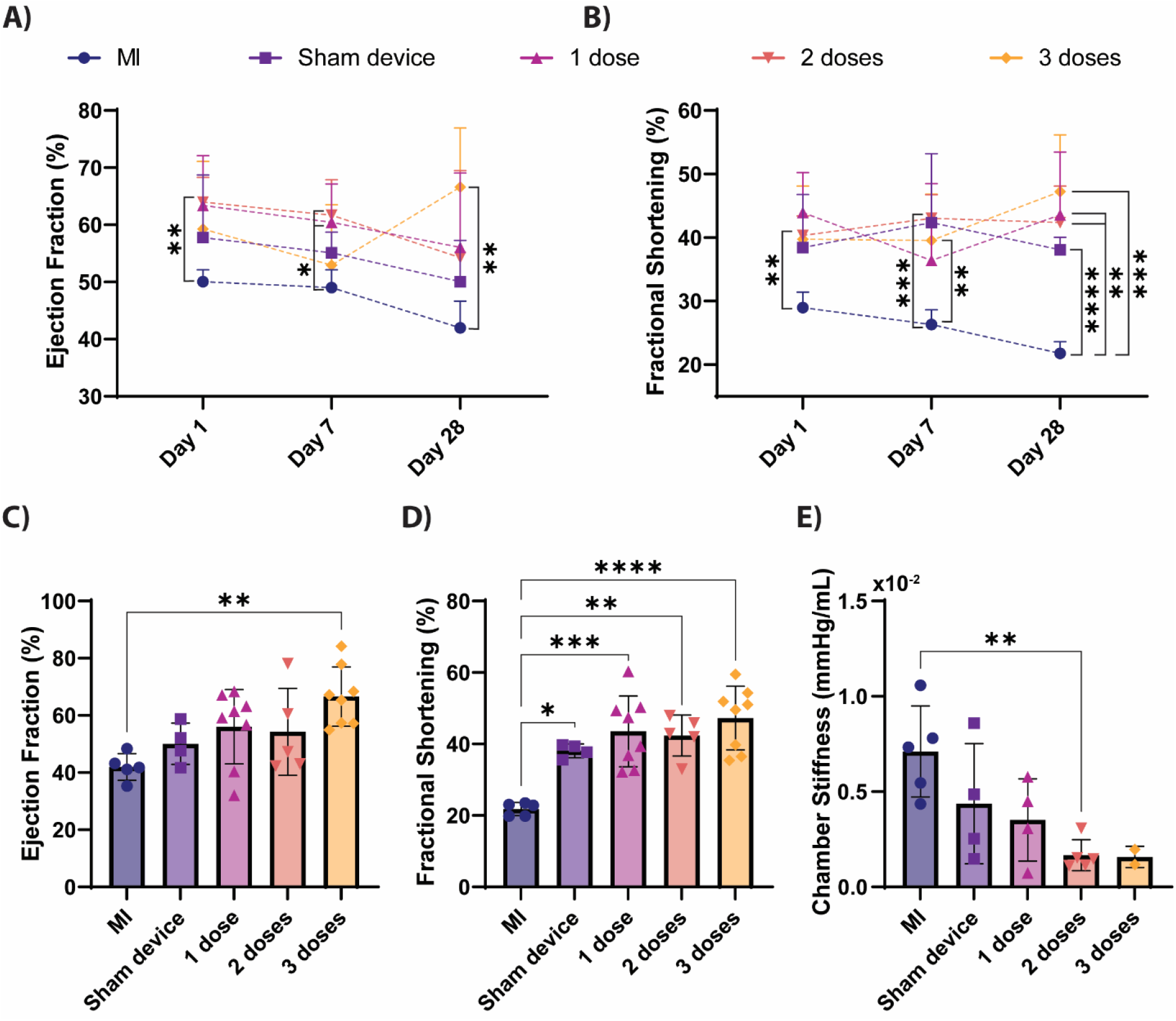
Cardiac function assessment. Ejection fraction. **(A)** and fractional shortening **(B)** for all groups at all study timepoints as assessed by echocardiography. Ejection fraction **(C)** and fractional shortening **(D)** at day 28 post-MI for all groups as assessed by echocardiography. **E)** Chamber stiffness at day 28 post-MI for all groups obtained from hemodynamic measurements. * P < 0.05, ** P < 0.005, *** P < 0.0005, ****P < 0.0001. Data are mean ± s.d. (n = 2-8) as analyzed by a one-way ANOVA (Mixed model) with Tukey’s multiple comparisons post-test. Individual animal values are shown as points in **C-E.**

By performing invasive hemodynamic measurements in a subset of animals 28 days post-MI, we show that left ventricular chamber stiffness decreases with FSTL1 delivery in a seemingly dose-dependent manner **(Figure 2e).** An FSTL1-driven reduction in infarct size and subsequent smaller scar region could explain this decrease in chamber stiffness as well as the recovery in EF and FS in the FSTL1-treated animals when compared to untreated controls. Importantly, chamber stiffness is a parameter derived from the EDPVR which reflects both myocardial mechanical properties and LV chamber geometry.^[25]^ Changes in EDPVR-derived parameters could arise from changes in myocardial properties or changes in LV geometry. Since we observed that the EDV (an indicator of LV geometry) did not significantly vary between our study groups **(Figure S1),** the improvement in chamber compliance we report could be attributed to reducing the region of the LV that stiffens due to scar formation through previously reported FSTL1-driven processes.^[17]^

Our results indicate that implantation of our epicardial device led to similar functional changes in the FSTL1-treated and sham device groups. In the absence of FSTL1, the functional changes in the sham device group likely arise from the mechanical benefit that implantation of the device itself has on the infarcted heart. Coupling biologic-free devices and/or biomaterials to the epicardial surface after MI can lead to functional benefits by changing the biomechanics of the infarcted LV through mechanical restraint and stress transfer to improve pump function or by limiting the degree of LV dilation and functional decrease over the course of MI remodeling.^[21,26–28]^ Our device is likely limiting the reduction in EF and FS over the course of post-MI remodeling by limiting the maximal degree of LV dilation that can occur.

We did not detect significant differences between the dosing regimens in terms of EF, FS or chamber stiffness were observed among the single, double, and triple dose regimens even though all functional parameters were better than the no treatment group. Some sources of noise potentially introducing variability into EF and FS quantifications include variations in MI size/location despite consistent position of LAD ligation (LV akinesis or thinning is not always visible at the mid-papillary plane used to calculate FS) and diminished rat echogenicity based on animal-specific anatomical features. Although a mouse model would have been more echogenic and has been used for other FSTL1 studies, miniaturization of the epicardial device platform to mouse size is challenging. A limitation of this study is that invasive hemodynamic recordings were only performed in only a subset of the animals. Future studies could further elucidate the physiological impact of multidose FSTL1 regimes by mitigating these sources of variability and limitations through MRI-based quantifications of cardiac function and biomechanics (e.g., strain). Monitoring clinically relevant biomarkers could allow for clustering of animals based on disease severity or treatment response and further elucidate if additional FSTL1 doses lead to a cumulative functional impact in certain disease conditions or is simply limited in our chosen model.

### 3.2. Multidose FSTL1 regimes lead to improved healing, remodeling, and angiogenesis

To further assess the FSTL1 cardiac benefits that may not be fully captured with echo-based functional metrics, we histologically assessed the impact of FSTL1 in infarct size, ventricular thickness, and angiogenesis in an arbitrary subset of excised heart samples.

#### Infarct size and LV thickness

Inspection of histological samples revealed prominent scar formation indicating a MI had occurred in all groups which is represented by hypochromatic staining in H&E sections and blue stained collagen in Masson’s trichrome sections **(Figure 3a).** A significant reduction of scarring **(Figure 3b)** was observed when 1, 2, or 3 doses of FSTL1 are delivered when compared to the MI only and sham device group. The double and triple FSTL1 dose regimens also led to a reduction of ventricular thinning compared to the MI only group **(Figure 3c).** Since neither the 1 dose nor the sham device group lead to a change in ventricular thickness, the double and triple dose regimens are likely mitigating the reduction of infarct extension post-MI solely biologically, via an FSTL1 specific pathway, or through a combined biomechanical mechanism, induced by FSTL1 and the mechanical benefit imparted by the epicardial device.

**Figure 3.**
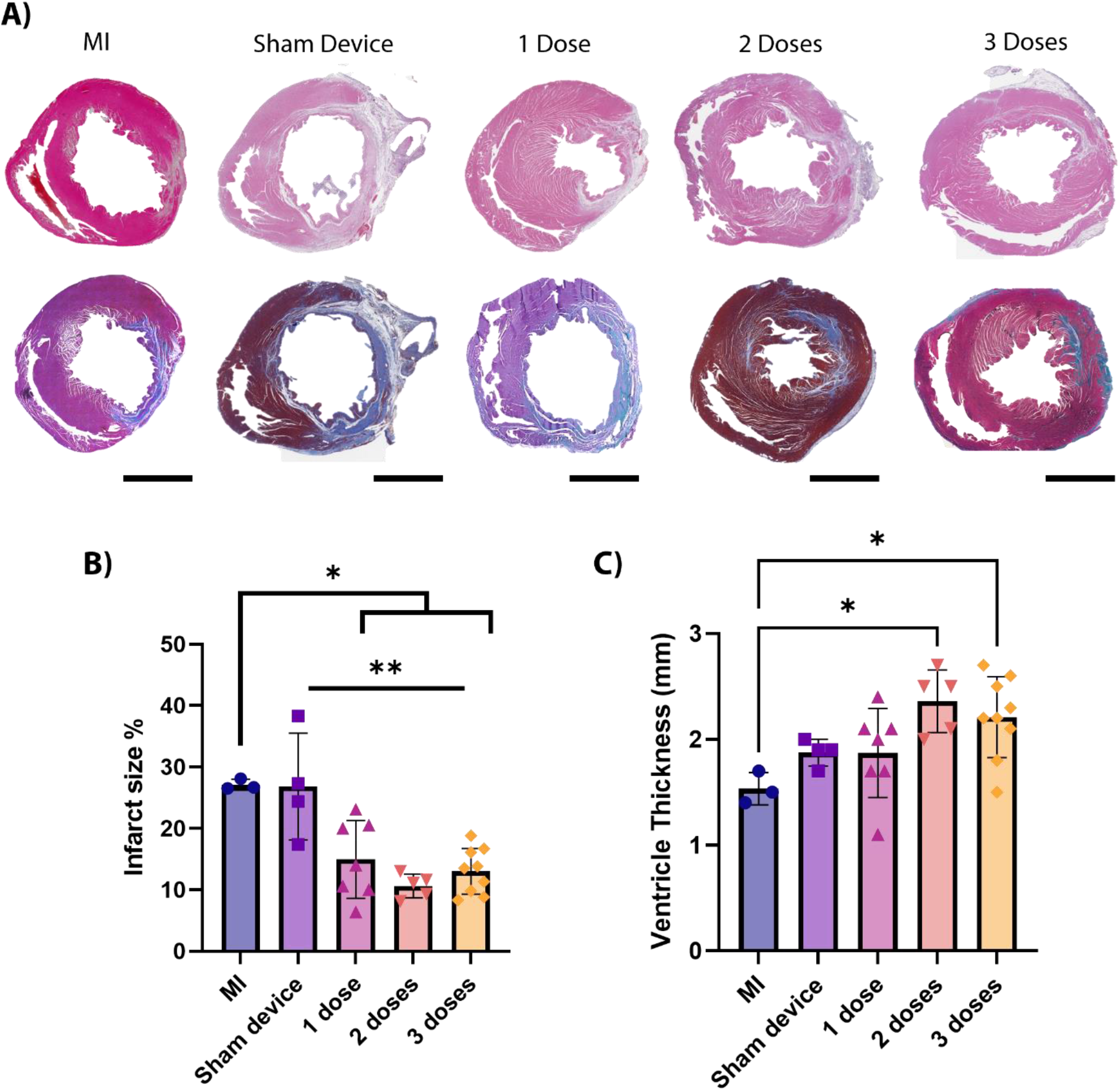
Histological assessment of infarct size and ventricle thickness. **A)** Representative H&E (top row) and Masson’s trichrome (bottom row) stained sections per group (scale bars are 2mm). Infarct size **(B)** and ventricular thickness **(C)** as quantified by histology. *P<0.05, **P<0.005. Data are mean ± s.d. as analyzed by a one-way ANOVA with Tukey’s multiple comparisons tests. Individual animal values are shown as points in **B-C**.

#### Angiogenesis

Three distinct zones were identified in all heart samples: the infarct zone (IZ) which was defined by a prominent collagen scar, the border zone (BZ), which surrounds the IZ, is composed of hypertrophic cardiomyocytes, and the adjacent myocardium (AM) composed of morphologically healthy cardiomyocytes with prominent striations. Representative CD31-stained sections from the IZ, BZ and AM were used to quantify blood vessel number and radial diffusion, a metric of vessel-vessel proximity **(Figure 4, Figure S2)**. As previously reported in a mouse model,^[17]^ we observed that exogenous FSTL1 induces a pro-angiogenic response in the IZ when compared to MI. Representative images of the IZ for all groups can be seen in **Figure 4a**. The triple FSTL1 dose regimen significantly increased the number of blood vessels **(Figure 4b)** and decreased neo-vessel-to-vessel distance (measured by radial diffusion, **Figure 4c**) in the IZ when compared to all control and treatment groups.

**Figure 4.**
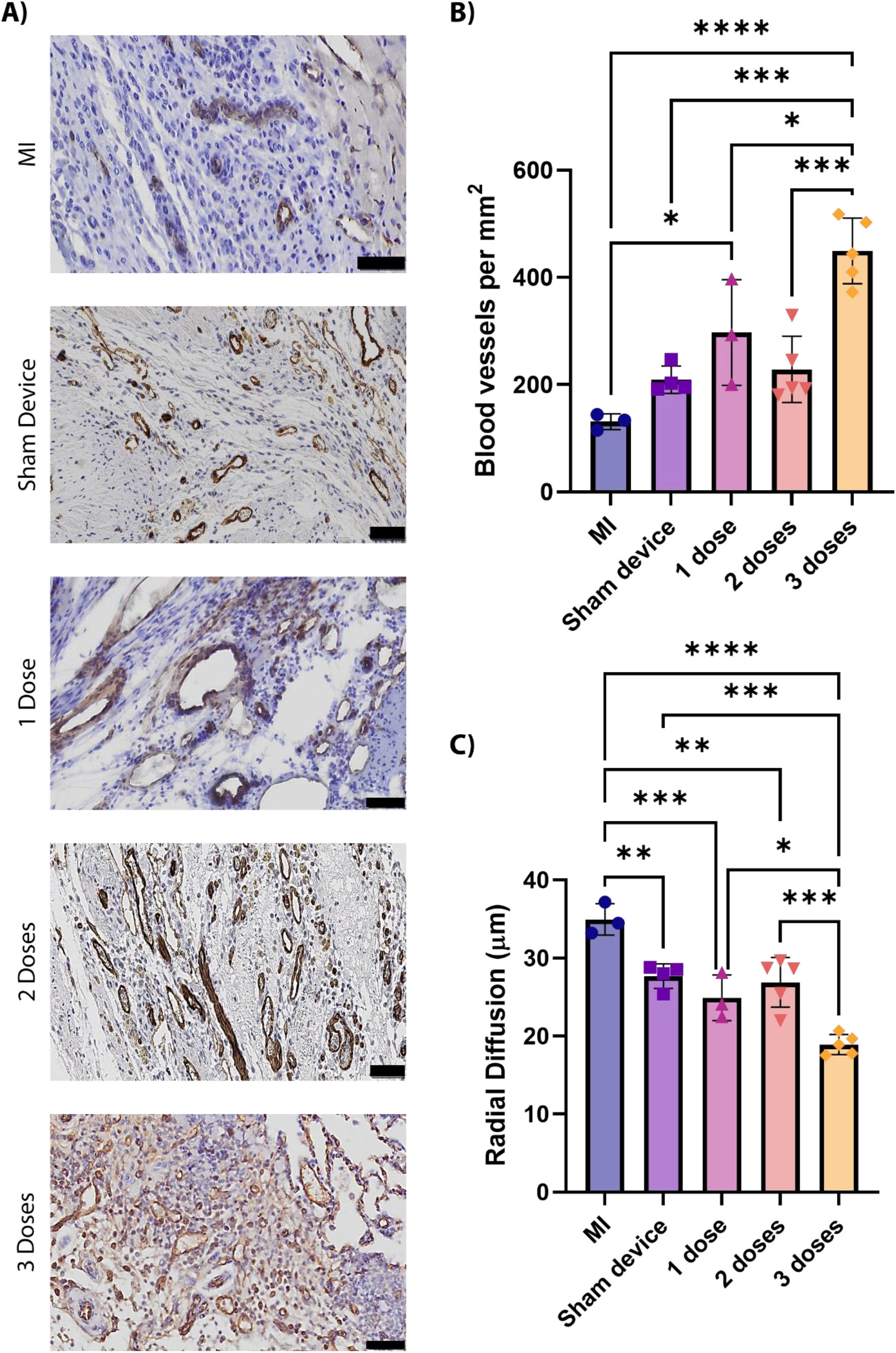
Histological assessment of angiogenesis. **A)** Representative CD31-stained sections per group (scale bar is 50 mm). Blood vessel number **(B)** and density **(C)** in the infarct zone as quantified by stereology. * P < 0.05, ** P < 0.005, *** P < 0.0005, ****P < 0.0001. Data are mean ± s.d. (n = 3–5) as analyzed by a one-way ANOVA (Mixed model) with Tukey’s multiple comparisons post-test. Individual animal values are shown as points in **B-C.**

An improvement in both blood vessel number and radial diffusion was seen with the single and double FSTL1 dose regimens when compared to the MI only group. However, this improvement is comparable to the improvement observed with the sham device group which, in the absence of FSTL1, could be attributed to the interaction between post-MI healing and the foreign body response to the gelatin biomaterial and implanted device. Further characterization of this interaction could be done by modulating the foreign body response to this indwelling device through the incorporation of mechanical actuation.^[29,30]^ Additionally, inclusion of a group that delivers a single dose of FSTL1 using our epicardial device, and not only a gelatin sponge, could help elucidate how differences in the foreign body response may influence this dosing regimen.

Improving the endogenous angiogenic response after MI can reduce scarring and adverse left ventricular remodeling.^[31]^ In this study, the triple FSTL1 dose regime proved to be superior at inducing a pro-angiogenic response when compared to both the single and double dose regimens, despite comparable functional improvements. The time of delivery of the 3rd FSTL1 dose may be important for generating an improvement in angiogenesis and is likely related to the interactions between FSTL1 and biological processes/cells characteristic of post-MI healing at day 21.^[31,32]^ Further studies are warranted to better understand and possibly further optimize the time-specific therapeutic effect driven by multiple FSTL1 dosing regimens.

The epicardial device used for this study enables multidose delivery of bioagents *in vivo* without additional invasive procedures and has broad utility outside of this specific example. This delivery platform can be used to optimize the dosing regimen of FSTL1 and assess other bioagents for MI treatment and applications in other pathologies. It can also enable multi-agent therapeutic regimens. Incorporating more advanced features to our epicardial delivery platform such as continuous sensing for closed loop bioagent delivery, may be required to maximize its therapeutic benefit for certain applications and increase its versatility which can ultimately contribute to its adoption in the clinic. This was the first study to deliver additional doses of FSTL1 in a non-invasive manner following implantation of an indwelling epicardial delivery platform in a rat model of MI.

## 4. Conclusions

In this study, we used an epicardial device to deliver multiple FSTL1 doses non-invasively and characterized the therapeutic benefit of single, double, and triple dose regimens without the effects of additional surgical interventions. We can draw the following conclusions from this study: 1) the beneficial effects of epicardial FSTL1 delivery previously reported in mouse models can be recapitulated with a rat model of MI and a gelatin sponge biomaterial carrier, 2) a single dose of FSTL1 yields either a higher EF and/or FS in FSTL1 treated groups in comparison to MI controls, 3) by day 28 after MI, 3 doses of FSTL1 lead to the greatest increase in EF and FS in comparison to MI controls, 4) FSTL1 decreases chamber stiffness after 2 doses and in seemingly dose-dependent manner, 5) a significant reduction in infarct size is observed after delivering 1-3 FSTL1 doses, 6) multidose FSTL1 regimens lead to increased ventricular wall thickness in comparison to MI controls, 7) FSTL1 increases blood vessel number and density in the infarct zone with 3 doses resulting in significantly improved angiogenesis in comparison to all other groups.

In summary, we report the first study where post-surgical doses of FSTL-1 could be delivered non-invasively with an implanted epicardial device. Multidosing results in improved cardiac function, healing, and angiogenesis. Our delivery platform may play an essential role in facile assessment of timed dosing regimens of different bioagent combinations for a range of clinical indications.

## Acknowledgements

The authors would like to thank: Kenneth Walsh, Vahid Serpooshan and Pilar Ruiz-Lozano for useful discussions during the planning stage of this study, Fred Roberts from FUJIFILM VisualSonics, Inc. for technical support with ultrasound, and members of the Hope Babette Tang (1983) Histology core for assistance with histological processing. C.E.V. acknowledges financial support from the Ford Foundation predoctoral fellowship and the NSF Graduate Research Fellowship Program. D.S.M. acknowledges financial support from the Irish Research Council Government of Ireland Postgraduate Scholarship (GOIPG/2017/927) and a Fulbright Enterprise Ireland Award. W.W. and G.P.D. acknowledge financial support from Science Foundation Ireland under grant SFI/12/RC/2278, Advanced Materials and Bioengineering Research (AMBER) Centre, University of Galway in Ireland. J.B. acknowledges funding from Lausanne University Hospital Improvement Fund and SICPA Foundation. E.T.R acknowledges departmental funding from the Institute for Medical Engineering and Science, the Mechanical Engineering Department at the Massachusetts Institute of Technology, and from NSF EFRI grant 1935291.

