## Supporting Information for "Epicardial reservoir-enabled multidose delivery of exogenous FSTL1 leads to improved cardiac function, healing, and angiogenesis"

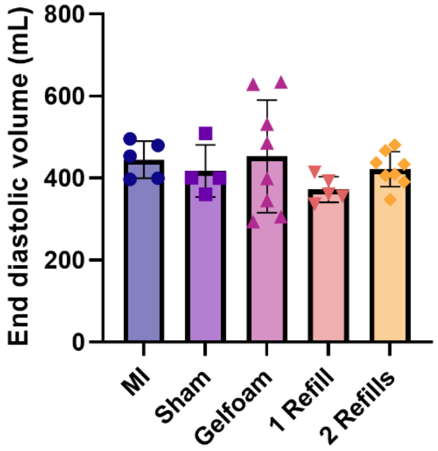


**Figure S1. End diastolic volume as assessed by echocardiography.** Data are mean ± s.d. (n =4–8) as analyzed by a one-way ANOVA (Mixed model) with Tukey's multiple comparisons post-test. Individual values are overlayed as points.

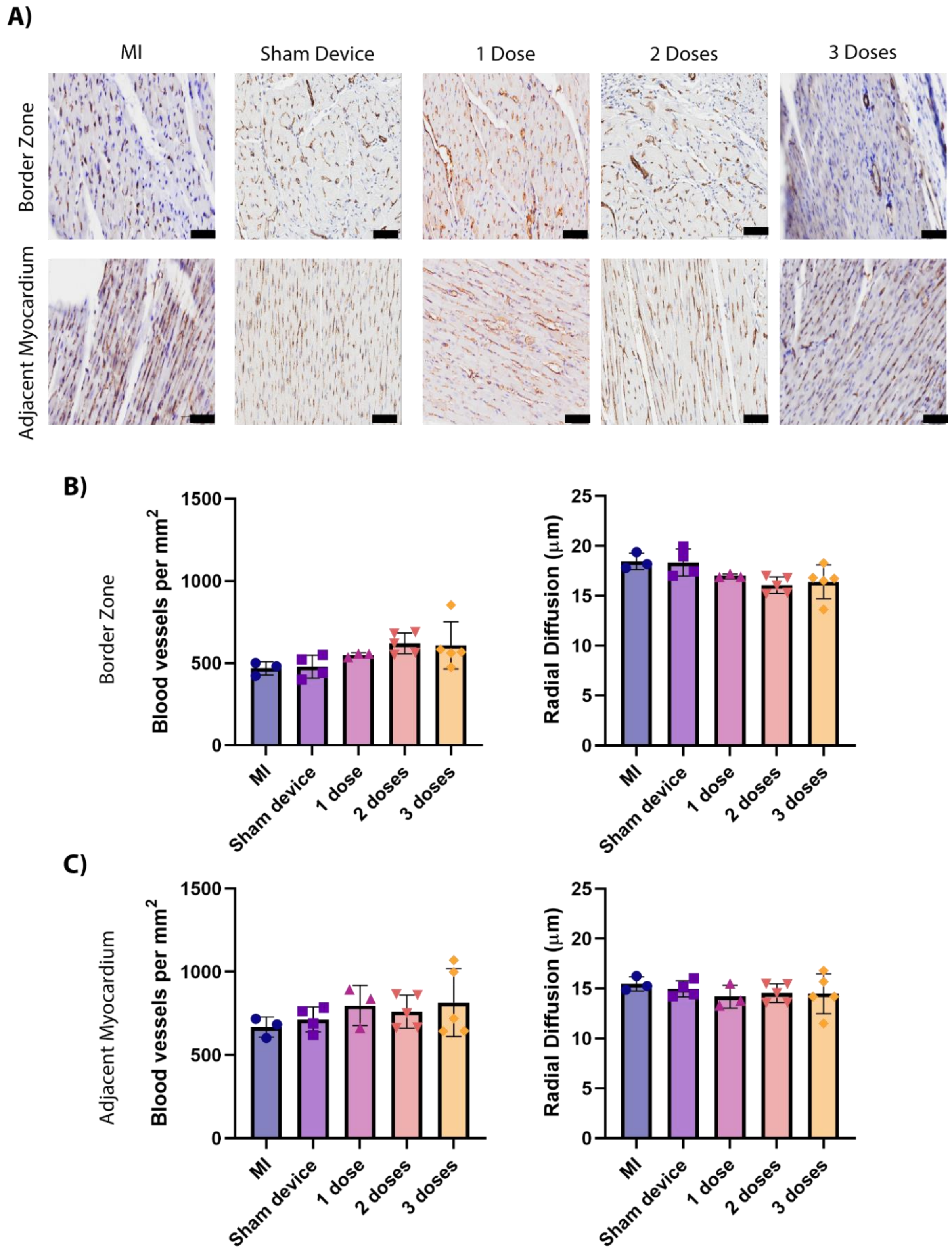


**<u>Figure S2. Additional histological assessment of angiogenesis.</u>** **A)** Representative CD31-stained sections of border zone and adjacent myocardium per group (scale bar is 50 mm). Blood vessel number and density in the border zone **(B)** and adjacent myocardium **(C)** as quantified by stereology. * $P < 0.05$, ** $P < 0.005$, *** $P < 0.0005$, ****$P < 0.0001$. Data are mean ± s.d. ($n = 3–5$) as analyzed by a one-way ANOVA (Mixed model) with Tukey's multiple comparisons post-test. Individual values are overlayed as points in **B-C.**
